# EVALUATION OF TAURINE’S NEUROPROTECTIVE EFFECTS ON SH-SY5Y CELLS UNDER OXIDATIVE STRESS

**DOI:** 10.1101/2021.04.26.441504

**Authors:** Rafaella Carvalho Rossato, Alessandro Eustaquio Campos Granato, Jessica Cristina Pinto, Carlos Dailton Guedes de Oliveira Moraes, Geisa Nogueira Salles, Cristina Pacheco Soares

## Abstract

Alzheimer’s disease (AD) is a type of dementia that affects millions of people. Although there is no cure, several study strategies seek to elucidate the mechanisms of the disease. Recent studies address the benefits of taurine. Thus, the present study aims to analyze the neuroprotective effect of taurine on human neuroblastoma, using an *in vitro* experimental model of oxidative stress induced by hydrocortisone in the SH-SY5Y cell line as a characteristic model of AD. The violet crystal assay was used for cell viability and the evaluation of cell morphology was performed by scanning electron microscopy (SEM). After pretreatment with taurine, the SH-SY5Y cell showed an improvement in cell viability in the face of oxidative stress and improved cell morphology. Thus, the treatment presented a neuroprotective effect.

**GRAPHICAL ABSTRACT:** 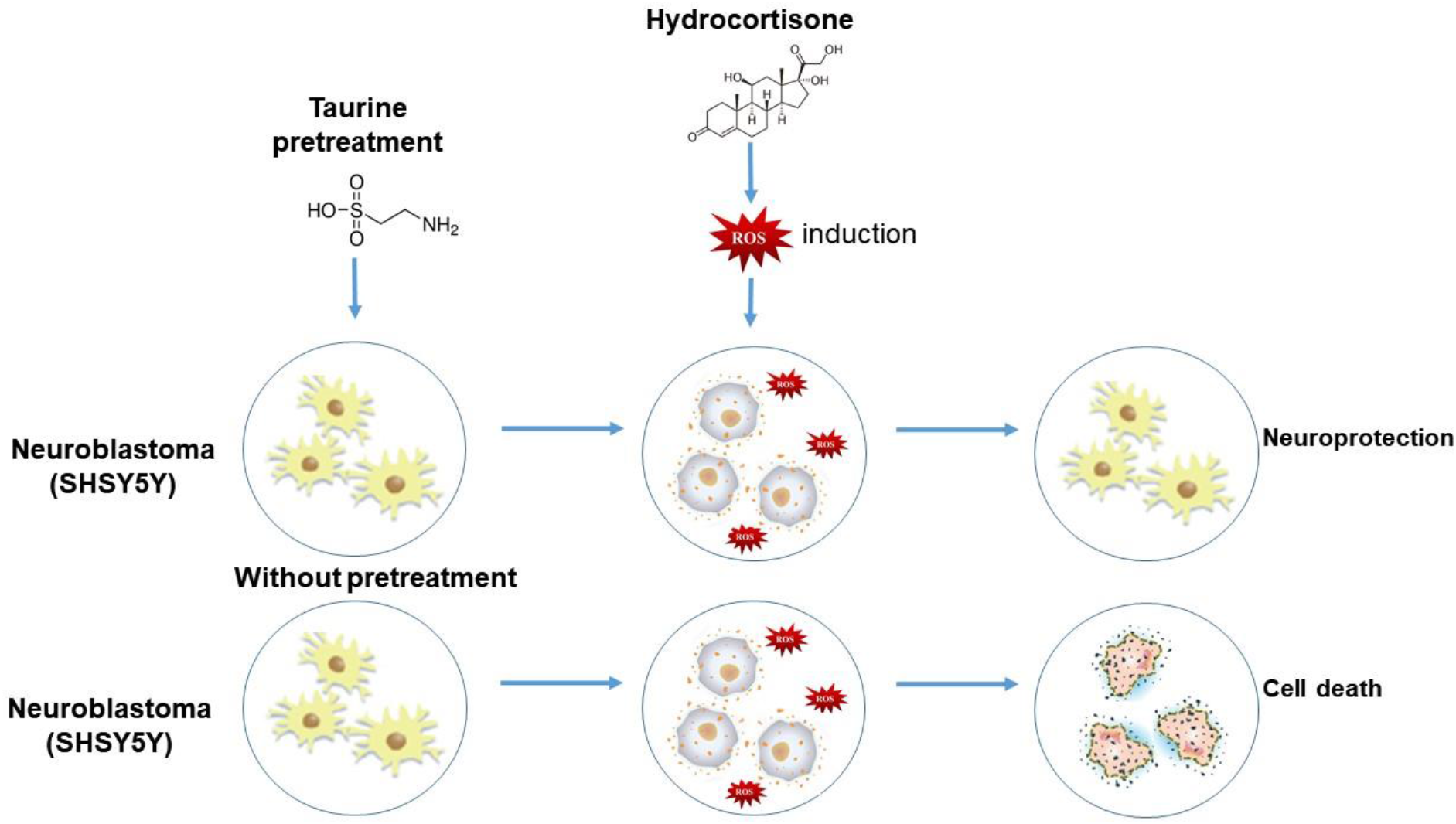

## INTRODUCTION

Alzheimer’s disease (AD) is the most common, progressive, and irreversible neurodegenerative disorder, characterized by memory loss, cognitive impairment, and behavioral abnormalities (REITZ; MAYEUX, 2014)(DE FALCO et al., 2016). The main risk factors are advanced age and family history; however, studies report that specific genetic mutations correlate with increased susceptibility, as well as the anticipation of clinical signs characteristic of AD (LIU et al., 2013) (REITZ; MAYEUX, 2014).

AD’s origin is still unknown, but some brain changes are described, including the abnormal production and deposit of β-amyloid protein and neurofibrillary tangles, resulted from TAU protein hyperphosphorylation (BELYAEV et al., 2010). The swedish mutation (SWE), represented as a specific modification in the gene that encodes the amyloid precursor protein (APP), can also be highlighted. Thus, the cleavage of APP β-secretase (codons 595 and 596 in APP695) makes APP become a preferable substrate for β-secretase (VIGNINI et al., 2011) (GIACOBINI; GOLD, 2013). As a result of this mutation, there is an acceleration of APP amyloidogenic processing. This fact is related to the secretion of exacerbated amounts of β- amyloid forms (Aβ) and causes the abnormal accumulation of intracellular Aβ (FERNANDES et al., 2018). Due to these biochemical changes caused by the altered genotype, the complexity of AD stands out.

The reduction in the number of nerve cells (neurons) and the connections between them (synapses) and other critical changes in the AD phenotype should be highlighted. Another point that can be studied is the degree of cognitive damage related to the dysfunction and specific degeneration of cholinergic neurons located in the basal forebrain’s cholinergic complex (BARBOSA, 2020). It is worth studying the cholinergic receptors that may represent one of the critical events in the pathology of this disease (GROTHE et al., 2014). As a consequence of the disease progression, there is a vast loss of neural prolongations, which leads to cerebral atrophy, decreases the weight of the brain mass, in addition to causing impairment in brain metabolism and regeneration capacity (NELSON et al., 2012).

In addition to genetic and biochemical changes, another probable reason for neural damage comes from oxidative stress obtained through the formation of reactive oxygen species (ROSs). They are unstable and extremely reactive molecules capable of altering other biomolecules such as proteins, carbohydrates, lipids, and nucleic acids. ROS are usually produced by body metabolism, but when they are produced in excess, they can exceed the cellular capacity for defense and repair, which leads to damage to neural cells (MORAES et al., 2019). It is evident that the chronic stress suffered by neural cells, associated with old age, predisposes to AD’s development. Psychological, behavioral, economic, and social risk factors can also be important as protagonists involved in cognitive decline and dementia (VITALIANO et al., 2011). These impairments in executive functions (attention and processing speed) and declarative memory, which is explicit memory evoked in the form of consciousness) are early clinical phenotypes for AD. The brain structures most sensitive to stress are the prefrontal cortex (PFC) and the hippocampus, and they are intricately linked to cognitive domains (SHANSKY, LIPPS; 2013). Those regions contain an exceptionally high density of receptors sensitive to cortisol, a hormone that is usually altered in acute and chronic stress (CORRÊA et al., 2016).

Hydrocortisone, a drug analogous to the hormone cortisol, can generate ROS, especially in high concentrations. Thus, it is worth mentioning the damage caused to biomolecules (ROSSATO et al, 2019). Neural cell’s normal functioning can be compromised when these damages are not repaired. Because of this, it provides cellular stress, which can evolve into atypical cells. Finally, the resolution of this process can lead, on the one hand, to cell death when this stress is not corrected; on the other hand, the neural cell survives; however, it becomes senescent and dysfunctional (SALLES et al., 2018) (MCEWEN, 2013). It is also possible to relate the changes in cortisol levels and ROSs with the decrease in brain-derived neurotrophic factor (BDNF), which modulate synaptic plasticity neurogenesis as neuronal help survival (JEANNETEAU; CHAO, 2013).

Counteracting chronic and oxidative stress is necessary. Taurine, or 2-aminoethanesulfonic acid, can be highlighted as promising for this function. Described as an agent capable of neuroprotection, it acts in functions of regulation of brain volume and maintains the integrity of the neural membrane and calcium homeostasis control. Thus, taurine can prevent the neural cell’s death since it can reduce oxidative stress (KILB; FUKUDA, 2017) (ZHOU et al., 2012). Taurine is found in abundance in the liver and brain, being synthesized from the metabolism of sulfur amino acids (methionine and cysteine), and is also considered a non-essential amino acid (MARCINKIEWICZ; KONTNY, 2014). In the central nervous system, taurine also functions as a chemical messenger. It is worth mentioning that the muscular system prevents oxidative stress in cells and, consequently, neuromuscular problems (WANG et al., 2013).

Therefore, the present study seeks to evaluate the effects of taurine in stress conditions with hydrocortisone in neural cells to simulate the metabolic and oxidative stress characteristic of AD. Thus, the study of neural cells’ behavior in vitro was performed by pretreatment of cells with taurine, which were later stressed with hydrocortisone, in order to study the neuroprotection mechanisms of taurine as a possible retarding agent of AD.

## MATERIALS AND METHODS

### Cell culture

SH-SY5Y cells (Human Neuroblastoma ATCC - CRL-2266) were cultivated in 25 cm^2^ cell culture bottles at 37 °C under 5 % CO2 in DMEM/F12 (Gibco - Dulbecco’s Modified Eagle Medium - Nutrient Mixture) supplemented with 10 % fetal bovine serum (Life Technologies) and 1 % antibiotic and antimycotic (penicillin and streptomycin - Thermo Fisher Scientific, Invitrogen) (SALLES et al., 2018).

### Cell plating

After the confluence grows in the culture bottles, the cells were detached with trypsin (0.01%) to carry out the experiments and cultivated in 96 well plates with a cellular density of 1 x 10^4^ cell/well for cell viability assay. For the analysis of cell morphology, SH-SY5Y cells were plated in 24 well plates on coverslips with a cellular density of 1 x 10^5^ cell/well.

### Experimental groups

The groups were described below according to the treatment the cells received.

**Control group:** cells not exposed to the stressor.

**Taurine only group:** cells exposed to taurine.

**Hydrocortisone only group:** cells exposed to hydrocortisone stressing agent.

**Taurine and hydrocortisone group:** cells exposed to taurine and hydrocortisone.

### Pretreatment

SH-SY5Y cells were pretreated with taurine (Sigma) (0.5 mg/mL) in DMEM/F12 (Gibco-Dulbecco’s Modified Eagle Medium - Nutrient Mixture) supplemented with 10 % fetal bovine serum (Life Technologies) and 1 % antibiotic and antimycotic (penicillin and streptomycin - Thermo Fisher Scientific, Invitrogen) (ROSSATO, 2019).

### Oxidative stress induction

SH-SY5Ycells were subjected to the hydrocortisone (Teuto) stressing agent at a concentration of 200 μM, dissolved in DMEM/F12 (Gibco - Dulbecco’s Modified Eagle Medium-Nutrient Mixture) supplemented with 10 % fetal bovine serum (Life Technologies) and 1 % antibiotic and antimycotic (penicillin and streptomycin - Thermo Fisher Scientific, Invitrogen) (ROSSATO, 2019).

### Cell viability assay

The cell viability assay evaluates the cytotoxicity of the compound employing a colorimetric test. The method consists of verifying the cell density by staining the DNA, obtaining quantitative information on the relative density of live cells adhered to culture plates, that is, obtaining indirectly the amount of DNA in the samples and consequently the number of cells present in the analyzed groups (MORAES et al., 2019).

The cells were washed with PBS and incubated in a 100 μL crystal violet solution (5%) for 4 min at room temperature and in the dark. The cells were then washed with running water to remove excess dye and incubated with Dimethyl sulfoxide (DMSO) for one hour. At the end of incubation, a SpectraCount - Packard 570 nm spectrophotometer reader was used for evaluated results. The whole process was carried out in the dark. The data collected were statistically analyzed (MORAES et al., 2019).

### Cell morphology analysis

The cells were washed three times with cacodylate buffer at a concentration of 0.1 M (SALLES et al., 2017) (SALLES et al., 2018) and fixed with the glutaraldehyde 2.5%, and the paraformaldehyde 4% in the cacodylate buffer 0.1 M and incubated for one hour. Next, dehydration was performed in a series of increasing acetone concentrations at concentrations of 30%, 50%, 70%, 90%, and 100%, for 10 minutes each pass. The acetone 100% + HMDS (Hexamethyldisilazane) in the proportion (1:1) for 10 minutes, and finally, with 100% HMDS for 10 minutes, was used for complete dry. The material was metalized in the EMITECH K 550 X® equipment, spraying a thin layer of gold on the samples. In this stage, the equipment metalized for 04 minutes and 50 seconds in a vacuum cycle using 17 kV. The cellular observation was performed using the Scanning Electron Microscope EVO MA10-Zeiss®.

### Statistical analysis

The present study was carried out with n = 8 and repeated three times separately to obtain confirmation. Statistical significance was admitted with p <0.05. GraphPad Prism 6® software was used, and the statistical tests used were the ANOVA ONE-WAY and the Tukey test.

#### Cell viability assay

**Figure 1:**
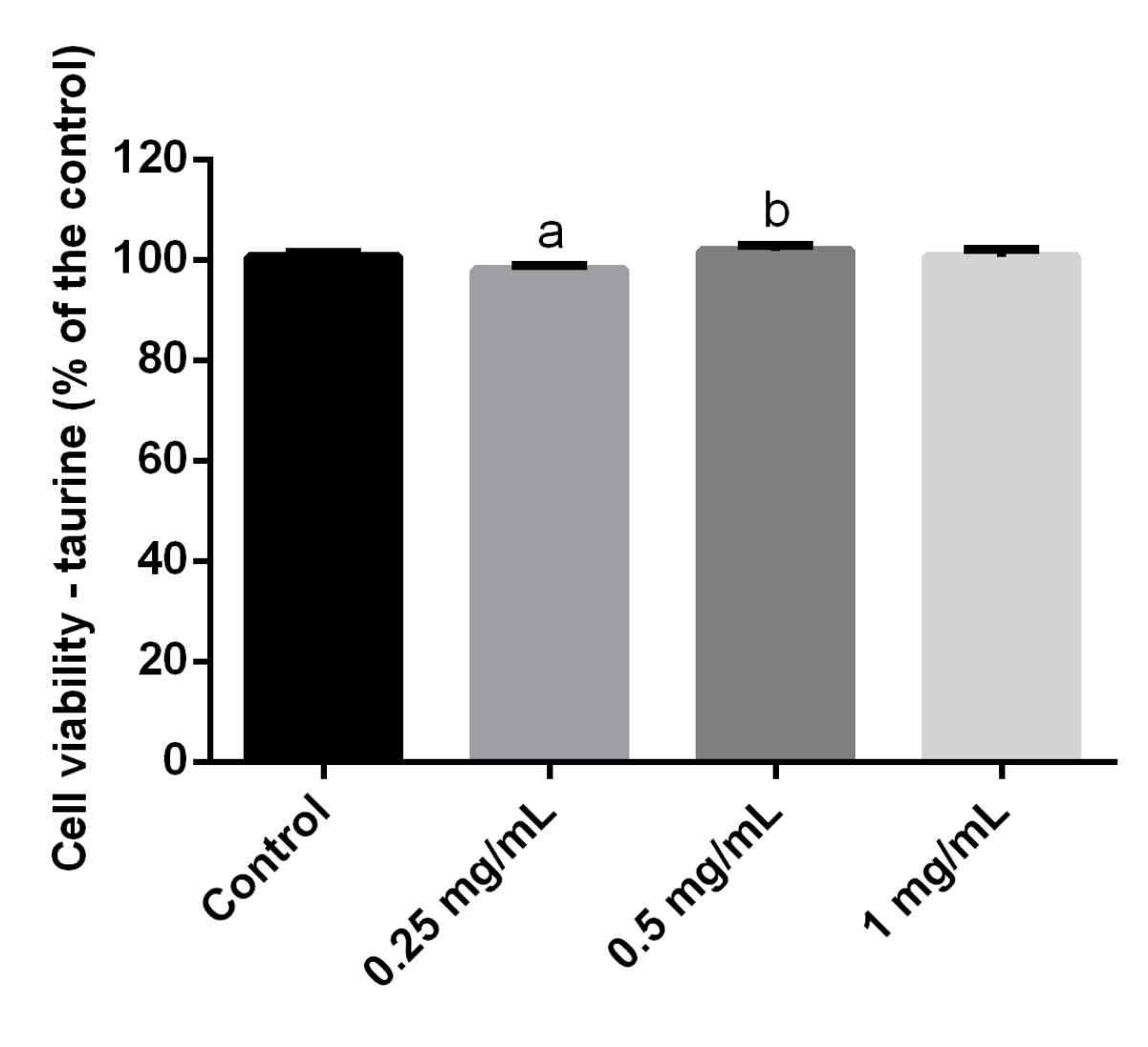
Percentage of cell viability on different concentrations of taurine (0.25; 0.5 and 1 mg/mL) compared to the control group.

The increasing concentrations of taurine showed that the 1 mg / mL concentration had similar behavior to the control group (without statistical difference). In contrast, the concentration of 0.25 mg / mL statistically decreased cell viability (p <0.0001) compared to the control group (a), and the concentration of 0.50 mg / mL showed a significant increase in cell viability (p = 0.0457) compared to control group (b).

**Figure 2:**
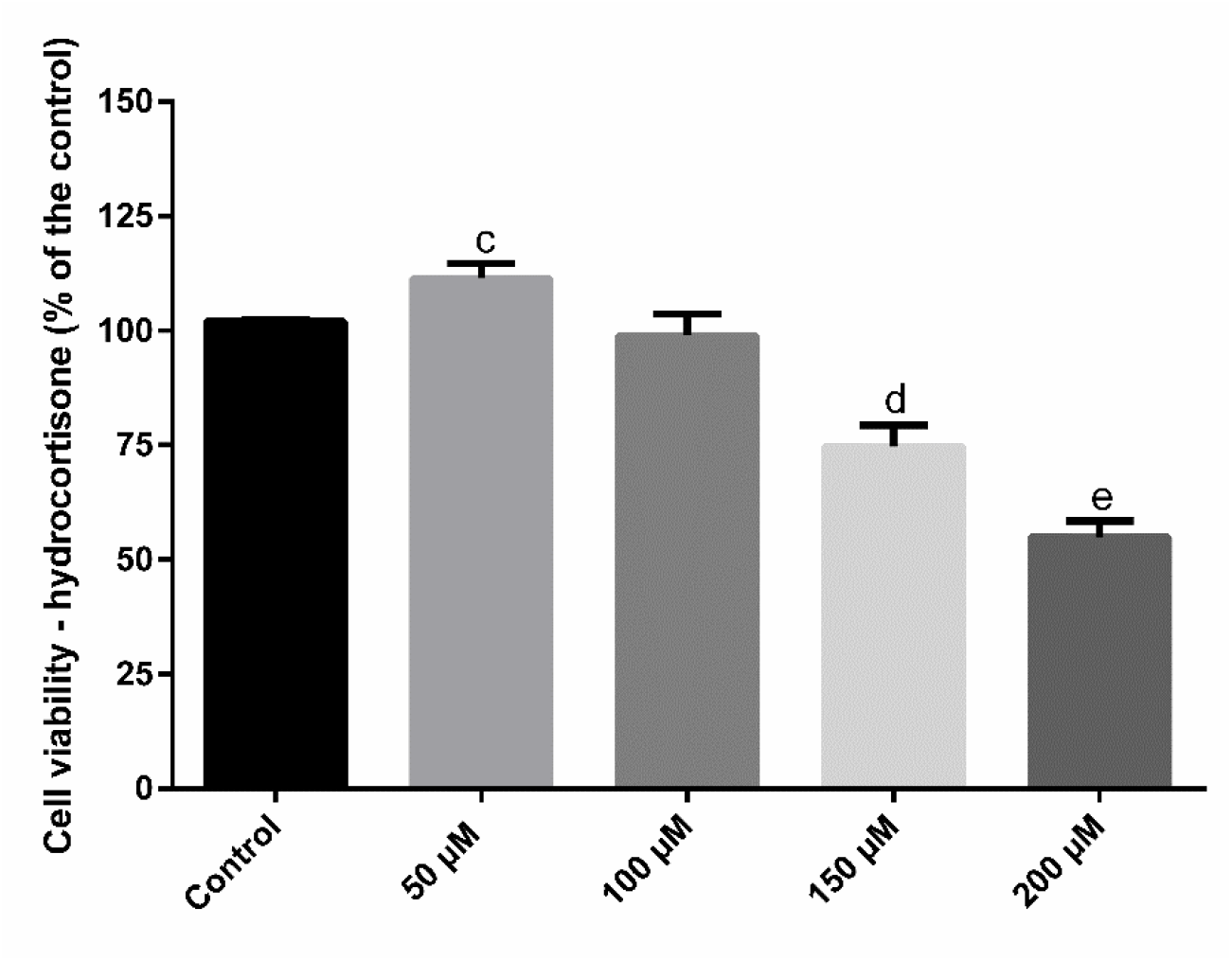
Percentage of cell viability on different concentrations of hydrocortisone (50; 100; 150 and 200 μM)

The increasing concentrations of hydrocortisone showed that the 100 μM had similar behavior when compared with the control group (without a statistical difference). On the other hand, the concentration of 50 μM statistically increased cell viability (p = 0.0051) compared to the control group (c), and concentrations of 150 and 200 μM showed a significant reduction in cell viability (p <0.0001), both compared to control group (d, e).

**Figure 3:**
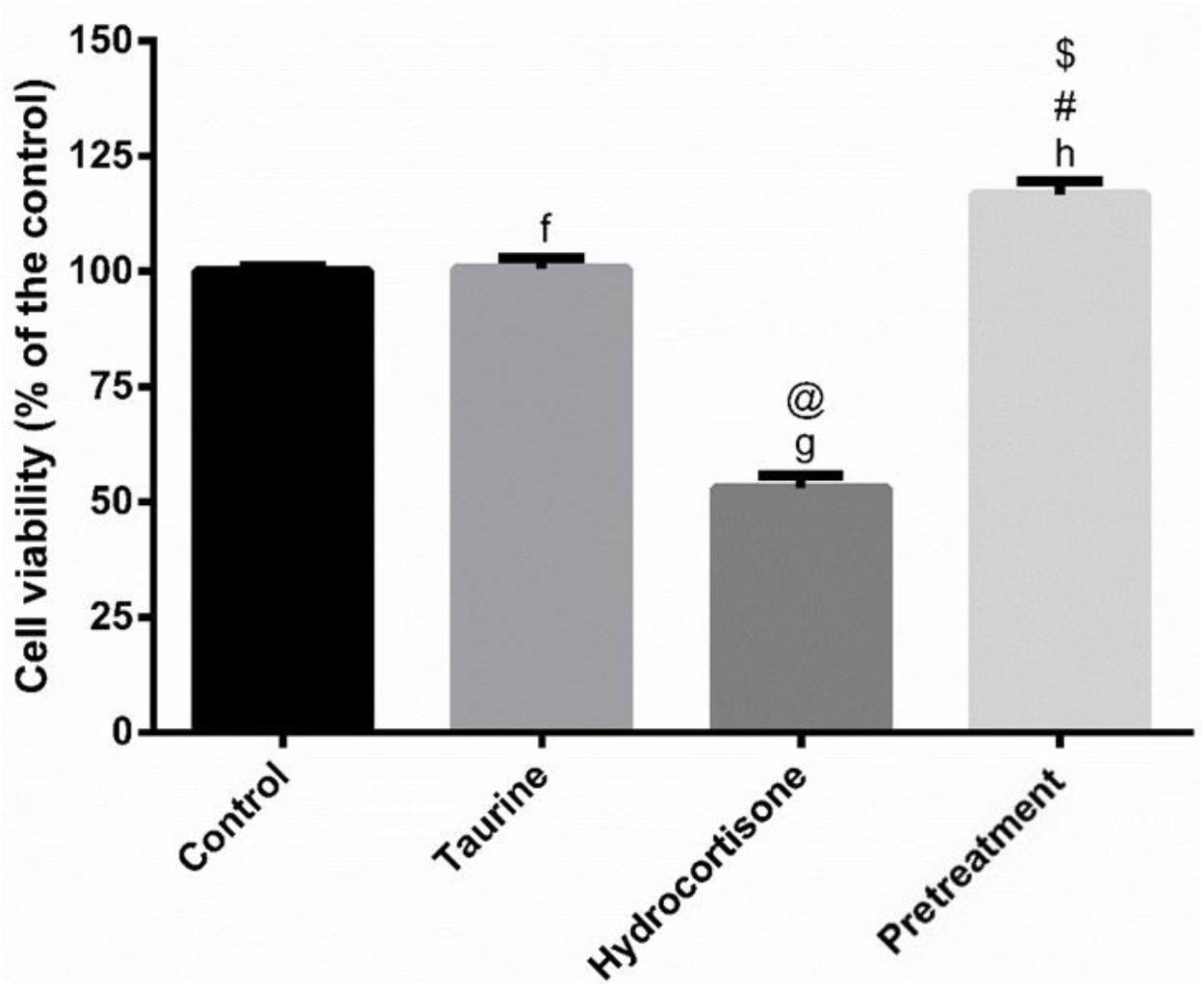
Percentage of cell viability on different concentrations of taurine (0.5 mg/mL), hydrocortisone (200 μM) and pretreatment.

When comparing cell viability, the taurine group (0.5 mg / mL) showed an increase in cell viability (p = 0.0457) compared to the control group (f). In the hydrocortisone group (200 μM), there was a decrease in cell viability (p <0.0001) compared to the control group (g), demonstrating the oxidative stress caused by this substance. The pretreatment group showed a significant increase in cell viability compared to the control group (h) (p <0.0001), demonstrating the anti-stress effect of Taurine. It is worth noting that the taurine group (0.5 mg / mL) also presents a statistical difference in relation to the hydrocortisone (@) (p <0.0001) and pretreatment (#) (p <0.0001) group. Finally, the pretreatment group showed a significant increase in cell viability concerning the hydrocortisone group ($) (p <0.0001).

#### Cell morphology analysis

**Figure 4:**
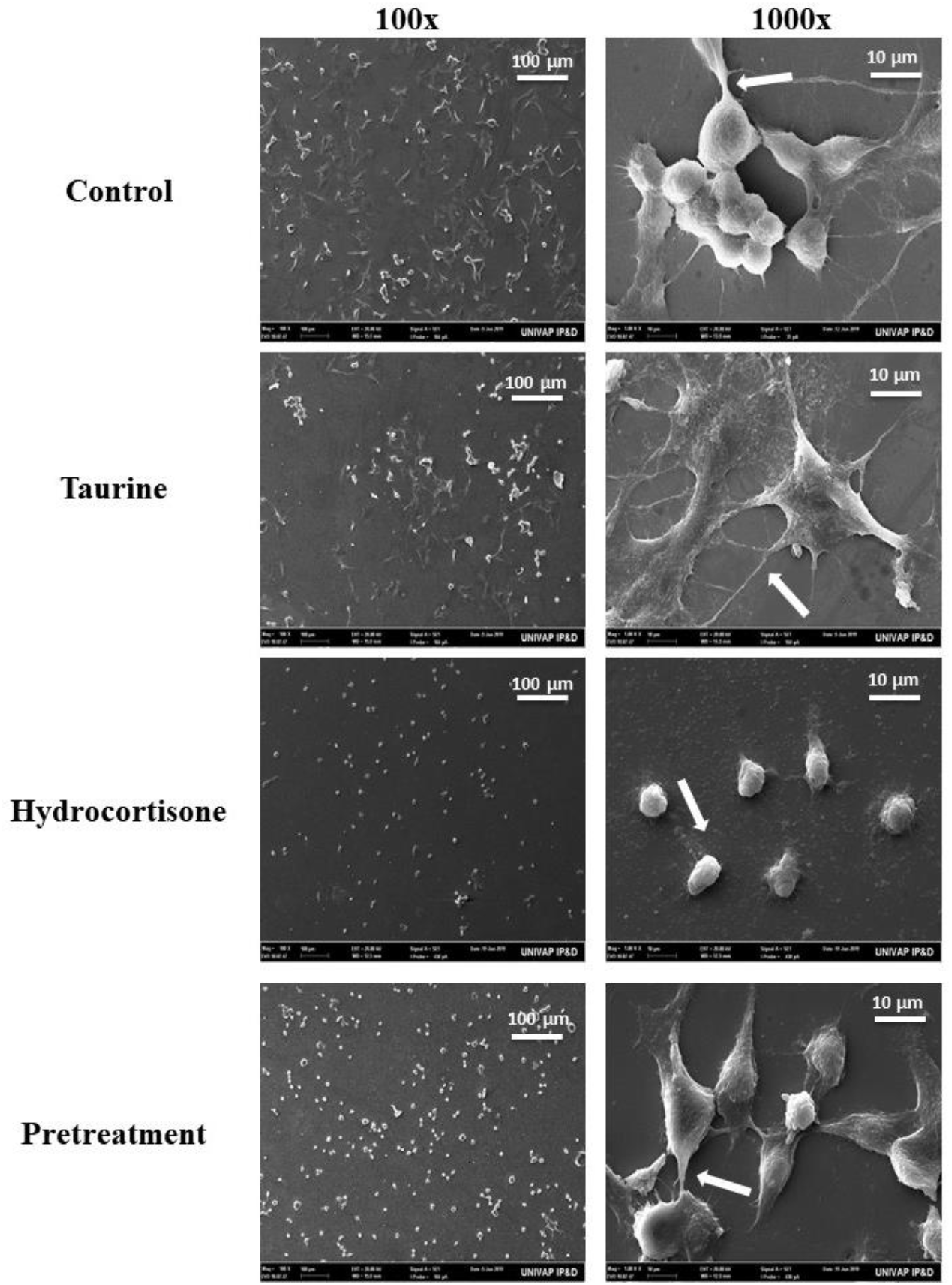
SEM images of the groups: Control, Taurine, Hydrocortisone and Pretreatment. The magnification in the first column is 100x and in the second 1000x.

Micrographs were obtained by scanning electron microscope (SEM) of SH-SY5Y cells. By analyzing the micrographs, we can verify the morphology of the SH-SY5Y cells. In the control group, it is possible to see the membrane projections characteristic of the neuroblastoma cell and the taurine group. In the stress groups with hydrocortisone, the cells lost their projections, presenting a retracted and rounded aspect. In the pretreatment group, a more significant number of cells and a recovery of cell morphology were observed.

**Figure 5:**
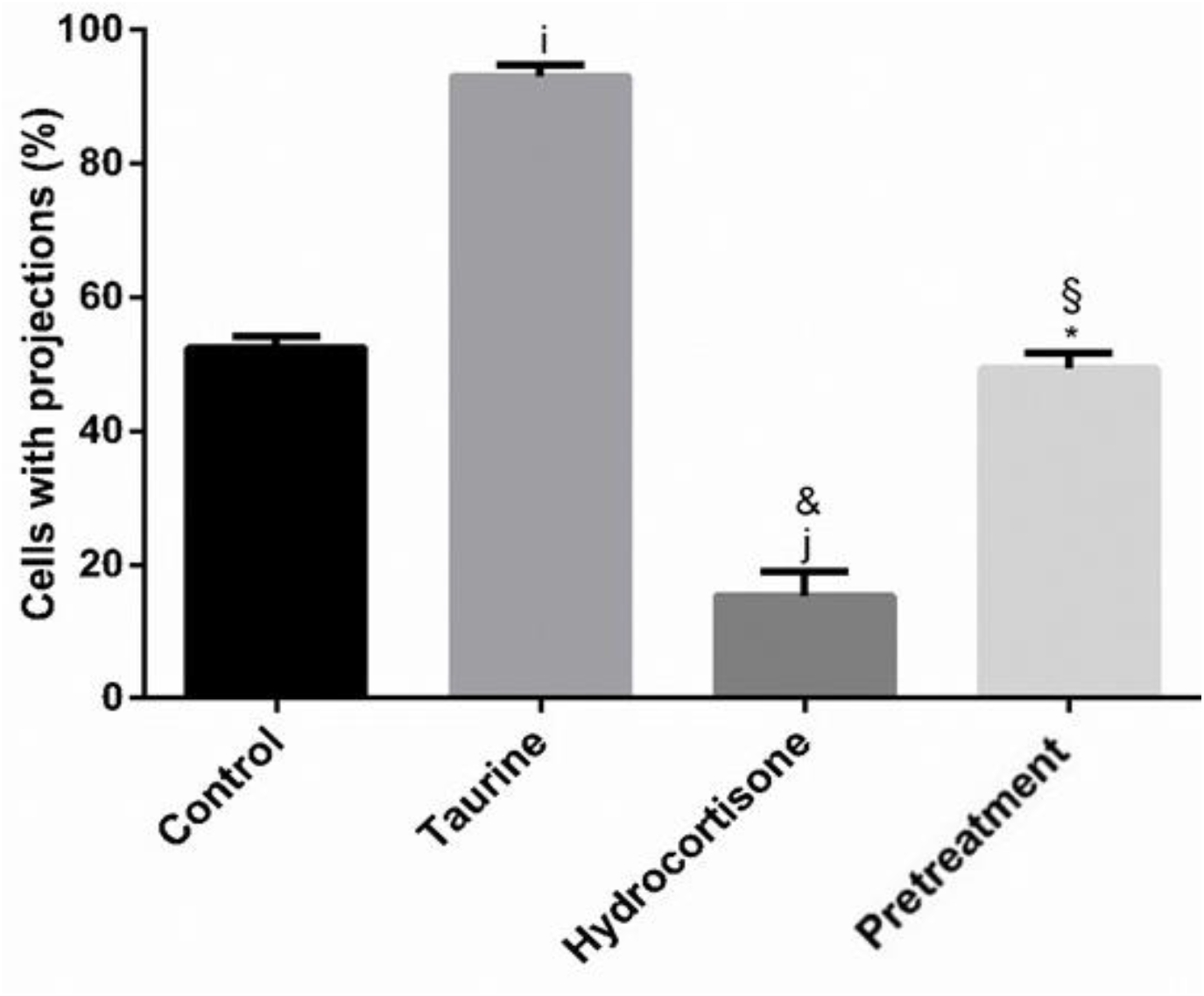
Percentage of cells with projections within the groups: Control, Taurine, Hydrocortisone and Treatment.

The frequency of prolongations in the neural cell line SH-SY5Y was counted using the micrographs obtained by scanning electron microscope (SEM). When analyzing, we can see an increase in the number of prolongations in the taurine group compared to the control (i) (p <0.0001). In the stress group with hydrocortisone, the cells showed a statistical decrease in prolongations compared to the control group (j) (p <0.0001). It is worth noting that the taurine group (0.5 mg / mL) also shows a statistical difference concerning the hydrocortisone (&) (p <0.0001) and pretreatment (*) (p <0.0001) group. Finally, the pretreatment group showed a significant increase in the frequency of neural cell extensions with the hydrocortisone group (§) (p <0.0001).

## DISCUSSION

This in vitro experimental investigation showed the capacity for metabolic and oxidative stress caused by hydrocortisone. Thus, this study establishes a potential experimental model for Alzheimer’s Disease to clarify the significant systemic effects, both in terms of the induction of systemic oxidative damage and the metabolic dysregulation provided by the hormone cortisol. Also, in this present study, the use of taurine is shown to be a promising alternative since it promoted neuroprotection in cells subjected to stress, as verified in the cell viability tests and scanning electron microscopy. Finally, it is possible to verify the interaction of taurine with neural cells in order to minimize the damage due to the metabolic and oxidative stress caused by hydrocortisone.

First, when conducting the study in the hydrocortisone group (a drug with a function similar to the hormone cortisol), it is possible to observe the cellular response of a dose-dependent character, that is, in lower doses and close to the standard threshold of the human organism, it presents an increase in cell viability. This event agrees with FONSECA; DA SILVA; SOARES, 2019, which establishes the positive effect on the organism in low concentration since it promotes glucose production maintenance.. LEE et al., 2017 affirms the relationship of cortisol in promoting cell protection in specific concentrations to improve cell viability. At the same time, a higher concentration of this hormone is possible to cause a reduction in cell viability, which is a fact evidenced in the study by DE LA RUBIA ORTÍ et al., 2017 with neural and brain damage and a reduction in the synapse in nervous tissue. TOLEDO et al., 2012 observed that high levels of cortisol alter the composition of cerebrospinal fluid, along with experimental evidence of damage to the hippocampus. Besides, studies by CURTO et al., 2017 highlight that increased plasma cortisol levels can be associated with the progression of cognitive deficit. The metabolic stress, related to high concentrations of cortisol, causes oxidative stress, thus enabling the appearance and acceleration of the clinical picture of neurodegenerative diseases.

To reverse the consequences of metabolic and oxidative stress resulting from high cortisol concentrations, supplementation with taurine is interesting for the gain in cell viability and neuroprotection evidenced by the cell viability assay. According to SARTORI, 2015, nutritional supplementation with taurine contributes to the immune system’s increased response since it acts on cellular osmoregulation and protein stabilization in cardiac and neural cells. A study by RIPPS; SHEN, 2012 highlights that taurine has an important neural role due to its cytoprotective capacity, which helps control salt concentrations, consequently improving cellular metabolic activity and antioxidation and cell development. Besides, a study by CONRADO et al., 2017 shows that taurine helps the central nervous system, thereby exercising a neurotransmitter function and improving the synapse. These results agree with PANDA; MISHRA; MISHRA, 2018 that reports the taurine ability to minimize oxidative stress, which acts in the control of salt concentrations in cells and the stabilization of the cell membrane.

The metabolic and oxidative stress effect of hydrocortisone was confirmed morphologically by SEM, which corroborates the results of cell viability. It is also possible to emphasize when observing through photomicrographs the characteristics and the morphological behavior of each cell. The images were obtained at a 100-fold increase, making it possible to view the cell population and a 1000-fold increase to check the cell morphology in each group. In this way, it is possible to observe in the hydrocortisone group that there was a structural change in the cytoskeleton, which retracted a neural cell’s characteristic prolongations. INELIA MORALES; GONZALO FAŔAS; RICARDO, 2010 relates these structural changes to oxidative stress, which states that ROSs causes metabolic stress. This relationship is positively harmful because when ROS is produced in excess in our body, it can lead to structural changes in proteins, lipids, and nucleic acids, leading to cell death. Studies by BAKER-NIGH et al., 2015 highlight the effect of oxidative stress on the cholinergic system, a fact presented as specific characteristics and vital importance for AD’s evolution. Since cholinergic neurons are particularly vulnerable to oxidative and metabolic stress, this causes damage to cholinergic signaling and, consequently, changes in neural morphology and its mechanisms.

WEI; JI, 2018 observed a reduction in the number of nerve cells and the connections between them, which can be correlated with the progressive reduction in brain volume confirmed by the study by DE FALCO et al., 2016 that verified the relationship between oxidative stress and the proliferation of astrocytes and activation of microglia, in order to justify the inflammatory character of AD. Together with the metabolic stress caused by cortisol, these changes can aggravate AD’s chronicity. Already SALAMEH et al., 2015 observed glycemic oscillations, can cause insulin resistance and metabolic alteration, as well as changes in the function of the insulin receptor and changes in the pattern of the vascular structure about the endothelial cells, in addition to an increase in astrocytes, cells which make up the BHE. According to RUIZ et al., 2016 causes signaling changes and in the growth factor that keep nerve cells healthy, which can disrupt glucose metabolism. In addition to being related to changes in the metabolism of beta-amyloid protein, as a result, it presents evidence of the close relationship between the pathways of the brain insulin receptors with the accumulation of beta-amyloid protein and the abnormal and hyperphosphorylated tau protein, oxidative stress, and metabolic. Our results agree with SALLES et al., 2018, which showed that oxidative stress causes morphological changes, loss of interaction and neural function, critical cellular events that can trigger cell death signaling. The study of LEE et al., 2017, corroborates the observation of neural failure in AD, thereby establishing a relationship between metabolic and oxidative stress caused in neural cells.

At the same time, it is possible to verify the neuroprotective effect of taurine by SEM, robust with cell viability results morphologically. Thus, it can be observed that taurine preserved the cytoskeleton and maintains the characteristic neural cell extensions and the spatial structure. According to TYAGI et al., 2015, this may be related to supplementation, thus increasing the expression of BDNF, thereby improving the function of the ion channel receptor, production of secondary messengers, and enzymatic activity, signal transmission, gene expression, and secretion of neurotransmitters. This way, it results in neural remodeling, which improved cell morphology since it maintained the neural cells’ projections ramifications.

Consequently, the neural cell shows an improvement in the response against ROSs, confirmed by RIPPS; SHEN, 2012, that emphasize taurine’s thrilling action as a promoter of cellular homeostasis and its role in improving mitochondrial function, which is in agreement with PANDA; MISHRA; MISHRA, 2018. According to SHIMADA, K., 2015 the anti-inflammatory character of taurine can help the nervous system and attenuate cellular senescence. The cytoprotective action of taurine is reaffirmed, according to WANG et al., 2018, which leads to the metabolic improvement of the neural cell, as well as helping in the development of DNA and preventing cell apoptosis. This occurs, according to Freitas, 2016 due to the ability of Taurine to facilitate cell permeability, thereby preventing formation and damage resulting from ROSs, in addition to functioning as a buffer that maintains the pH of the medium and captures electrically viable ions.

However, according to CORRÊA et al., 2015 cortisol is a steroid-related to metabolic and cellular stress. Thus, CORRÊA et al., 2016 suggest that cortisol is a reliable marker for cognitive changes. A study by ZVĚŘOVÁ et al., 2013 showed that cortisol increases are related to their patients’ cognitive decline. WANG et al., 2018 reports the improved response capacity of the hypothalamic adrenocortical axis (HPA) to noradrenergic stimulatory regulation in AD, as well as the rupture of the blood-brain barrier, contribute to the clinical picture since the noradrenergic stimulatory regulation of the brainstem of the HPA axis, are substantially increased in AD.

The analysis of neurites with high hydrocortisone concentrations provides great oxidative and, consequently, metabolic stress to the neural cell. According to REITZ; MAYEUX, 2014 the reduction in neurites’ density accompanies the decrease in cell viability in a dose-dependent manner to oxidative and consequently metabolic stress. Corroborates with JACK; HOLTZMAN, 2013, which indicates that hyperphosphorylation of tau and its subsequent deposition is related to the degeneration of neurons in AD patients’ brains and the decrease in prolongations which can be used as an essential biomarker for this disease pathology. According to KAWAHARA, 2012, it relates to the beta-amyloid aggregates, thereby showing the calcium channels’ responsibility in the cell membrane and allowing the ion to enter. These events can destabilize homeostasis and modify the characteristic morphology of the neural cell. These biochemical and morphological changes can culminate in cell death.

The supplementation with taurine reversed this scenario, in agreement with a study by HANSEN et al., 2010 the neural differentiation of SH-SY5Y cells and phenotypic changes, such as emission of cytoplasmic projections, reduction of proliferation, expression of dopaminergic markers. According to GNEGY, 2012 all this may be related to stimulation of tropomyosin kinase B’s expression (TrkB) receptor, which makes cells responsive to the brain-derived neurotrophic factor (BDNF), thus serving as a growth factor for neural cells. The differentiation of mature neuronal phenotypes under the right conditions is emphasized, resulting in more extensions and branched neurites.

## CONCLUSION

This work as a whole showed for the first time that taurine can promote neuroprotection in SH-SY5Y neural cells, which minimized the effects of the hydrocortisone metabolic stresses that were provided. Therefore, taurine can be a model of study for cases of neurodegenerative diseases such as Alzheimer’s disease.

## Notes

### Competing Interest Statement

The authors have declared no competing interest.

